# Podoplanin drives dedifferentiation and amoeboid invasion of melanoma

**DOI:** 10.1101/2020.07.23.218578

**Authors:** Charlotte M. de Winde, Samantha L. George, Abbey B. Arp, Agnesska C. Benjamin, Eva Crosas-Molist, Yukti Hari-Gupta, Alexander Carver, Valerio Imperatore, Victor G. Martinez, Victoria Sanz-Moreno, Sophie E. Acton

## Abstract

Melanoma is an aggressive skin cancer developing from melanocytes, frequently resulting in metastatic disease. Melanoma cells utilise amoeboid migration as mode of local invasion. Amoeboid invasion is characterized by rounded cell morphology and high actomyosin contractility driven by the RhoA signalling pathway. Migrastatic drugs targeting actin polymerization and contractility to inhibit invasion and metastasis are therefore a promising treatment option. To predict amoeboid invasion and metastatic potential, there is a need for biomarkers functionally linked to contractility pathways. The glycoprotein podoplanin drives actomyosin contractility in lymphoid fibroblasts, and is overexpressed in several cancer types. Here, we show that podoplanin enhances amoeboid invasion in melanoma. Expression of podoplanin in murine melanoma models drives rounded cell morphology, increasing motility and invasion *in vivo*. Podoplanin expression is upregulated in a subset of dedifferentiated human melanoma, and *in vitro* is sufficient to suppress melanogenesis and upregulate melanoma-associated markers *Mitf* and *Pou3f2*. Together, our data indicates that podoplanin is both a potential biomarker for dedifferentiated invasive melanoma and a promising migrastatic therapeutic target.

## Introduction

Metastatic melanoma has a very poor prognosis, and there is a need for additional treatment options. Mutation of *BRAF* (V600E) causes hyperactivation of proliferation and survival signalling pathways (MAPK, ERK) driving melanoma progression (1). Combinations of BRAF inhibitors and immunotherapies targeting PD-1/PD-L1 show encouraging results, however some patients develop resistance to treatment (2). A novel alternative treatment strategy is to directly inhibit metastatic spread with migrastatic drugs, inhibiting actin polymerisation and contractility mechanisms (3).

Metastasis is a multistep process, starting with local invasion of the surrounding tumour microenvironment, and leading to systemic spread (4). Cancer cells utilise, and can switch between, different modes of migration to invade through extracellular matrix and migrate to secondary sites (5). Amoeboid migration is rapid, and characterised by high actomyosin contractility driven by RhoA, Rho kinase (ROCK) and Myosin II signalling. Amoeboid invasive cells exhibit rounded cell morphology and continuous formation of protruding membrane blebs; allowing the cell to squeeze through the extracellular matrix (ECM) (6), and reduce the requirement for proteolytic degradation of ECM (5). Indeed, matrix metalloprotease inhibitors are ineffective in blocking amoeboid invasion (7).

The transition to metastatic disease is a multi-faceted change in cancer cell state, and not simply the initiation of cancer cell migration. Indeed, acquisition of invasive motility by melanoma cells has also been linked to a return to a more stem-like phenotype or dedifferentiation (8–10), closer to a trafficking melanoblast than a mature melanocyte. Moreover, high ROCK-Myosin II contractile signalling also drives changes in secretory pathways (11). Amoeboid-driven secretion of interleukin-1 alpha (IL-1*α*) and activation of NF-*κβ* signalling results in macrophage differentiation which further promotes tumour growth and invasion (11).

The cytoskeletal signalling cascades driving actomyosin contractility are well understood. However, there is also a need to identify potential biomarkers linked to contractility pathways, thereby predicting amoeboid invasion and metastatic progression. The membrane glycoprotein podoplanin (also known as gp38, Aggrus, PA2.26, D2-40, T1*α*) drives actomyosin contractility in lymphoid fibroblasts (12), and is furthermore associated with myofibroblast phenotypes in inflammation and cancer (13). We hypothesised that podoplanin also drives cytoskeletal contractility in cancer cells. Results in this study show that podoplanin expression drives contractile amoeboid morphology in melanoma, triggering invasion and promoting metastasis.

## Results

### Podoplanin expression is increased in human melanoma

Melanoma often acquires an amoeboid mode of invasion to metastasise, but the drivers of this conversion are incompletely understood. Podoplanin is known to be upregulated in many cancer types (13) and drives actomyosin contractility through the activation of RhoA GTPase in fibroblasts (12). Since amoeboid cell migration and invasion requires high actomyosin contractility (5), we asked whether podoplanin expression in melanoma could drive amoeboid invasion and promote metastasis. We examined podoplanin (gene *PDPN*) expression in the metastatic human melanoma cell line WM938B and its non-metastatic counterpart WM938A (14) (**Fig. 1A-C**), and across clinical datasets (**Fig. 1D**). WM983B cells acquire amoeboid morphology when cultured on top of deformable collagen-based matrices exhibiting a rounded, contracted phenotype (70%), whereas WM983A cells exhibit a more spread, mesenchymal phenotype (**Fig. 1A-B**). This shift to contracted, rounded morphology correlates with a 2.5-fold higher expression of *PDPN* mRNA (**Fig. 1C**), suggesting that metastatic amoeboid melanoma cells may have higher podoplanin expression. Examination of mRNA data in publicly available datasets from clinical primary melanoma biopsies confirmed that *PDPN* transcript levels are higher in melanoma samples than in controls (benign nevi or normal skin) (**Fig. 1D**).

**Fig. 1.**
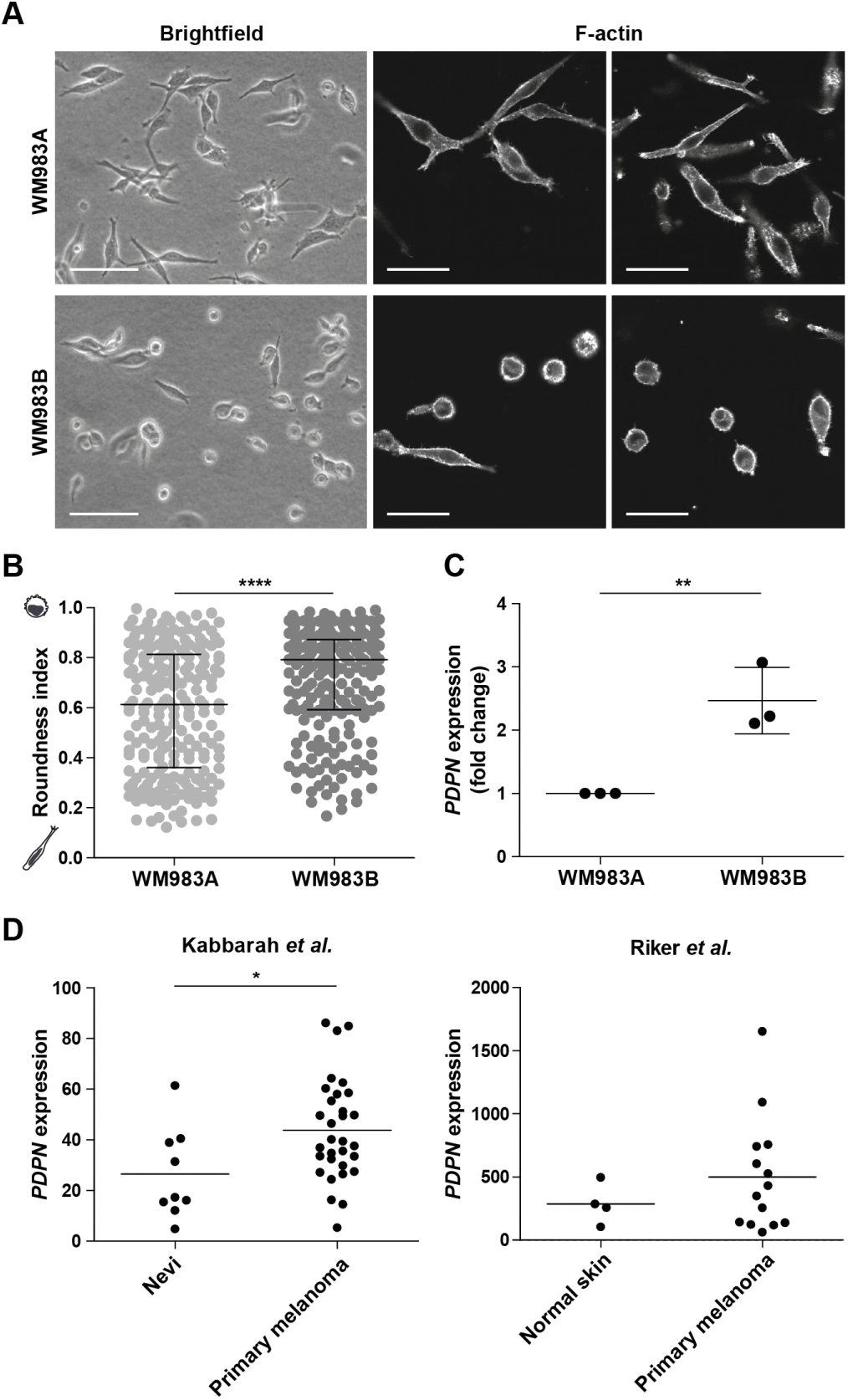
Increased podoplanin expression in human melanoma. **A**. Bright-field (left) and immunofluorescence (right) imaging of WM983A (top; primary) and WM983B (bottom; invasive/metastatic) human melanoma cell lines cultured on top of collagen matrix. Cells are stained for F-actin (white). Representative images from n=3 biological replicates are shown. The scale bars represent 45 (left) or 40 (right) microns. **B**. Cell morphology (roundness index) of WM983A (light grey) and WM983B (dark grey) cells cultured on top of collagen/Matrigel matrix. Data shown as median with interquartile range collated from 3-5 images per replicate of n=3 biological replicates. \*\*\*\**p*<0.0001. **C**. Podoplanin (*PDPN*) mRNA expression in WM983A and WM983B cell lines. mRNA expression is calculated as fold change and normalized to *GAPDH* expression. Data shown as mean +/-SD with dots representing n=3 biological replicates. \*\**p*=0.0084. **D**. Relative *PDPN* mRNA expression in two datasets (Kabbarah *et al*, GSE46517 and Riker *et al*, GSE7553) of human melanoma and appropriate control tissues. Data shown as mean with dots representing melanoma tumours from individual patients. \**p*=0.0250.

### Podoplanin promotes contractility in melanoma cells

We examined podoplanin expression in mouse models of melanoma and found that podoplanin surface protein expression is independent of *Braf* mutation status (**Fig. S1**). *Braf* mutant cell lines range from podoplanin low (5555) to high expression (4434). B16F10 melanoma cells (*Braf* -V600E mutation negative) express intermediate levels of podoplanin protein (**Fig. S1**). To directly test the contribution of podoplanin to melanoma morphology and invasion, we knocked out *Pdpn* (PDPN KO) in the metastatic cell line B16F10 using CRISPR/Cas9 technology. B16F10 cells predominantly express *Pdpn* transcript variant 1, similarly to lymph node stromal cells where endogenously high podoplanin levels drive cytoskeletal contractility (**Fig. 2A**). We confirmed that the guide RNAs successfully targeted podoplanin, and both *Pdpn* mRNA (**Fig. 2A**) and surface protein expression (**Fig. 2B**) were reduced to background levels. PDPN^+^ and PDPN KO B16F10 were then labelled with either mOrange or CFP for direct comparison in mixed cell cultures. In *in vitro* cell culture, PDPN^+^ cells grow in clusters with close cell to cell contacts (**Fig. 2C**). Each PDPN^+^ cell exhibits a rounded, contracted morphology, and intense F-actin staining is localised at the cell cortex (**Fig. 2C**). PDPN KO cells are dramatically more spread and exhibit multiple protrusions, as quantified by increased cell area and perimeter respectively (**Fig. 2C**), and no longer cluster. Additionally, PDPN KO B16F10 cells have lower F-actin intensity, and F-actin is arranged in filaments and puncta throughout the cell body (**Fig. 2C**).

**Fig. 2.**
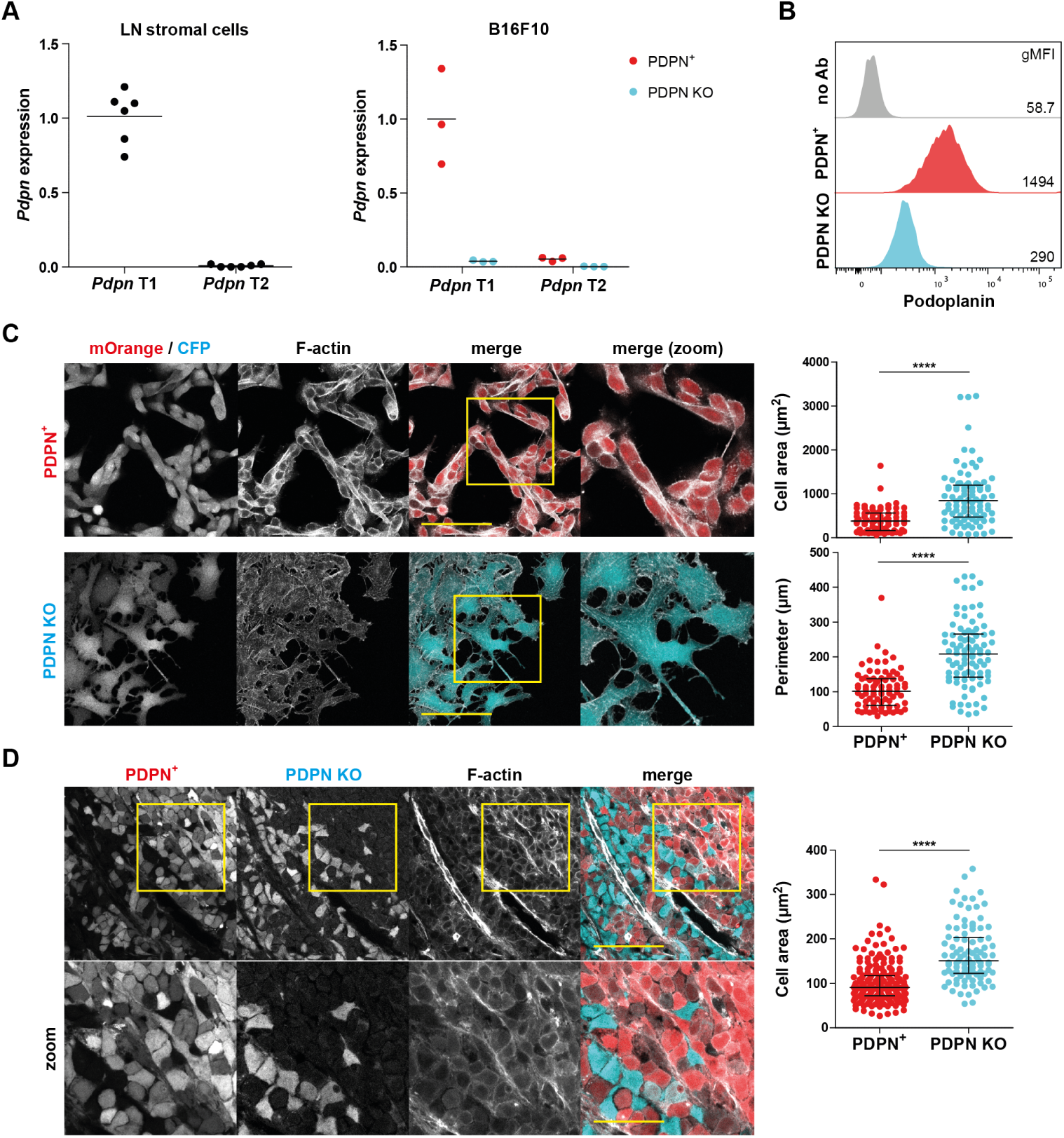
Podoplanin controls contractility of melanoma cells *in vitro* and *in vivo*. **A**. Expression of podoplanin (*Pdpn*) mRNA transcript variant 1 (T1) and 2 (T2) in lymph node (LN) stromal cells cultured *ex vivo* for 3 days (left), and PDPN^+^ (red) and podoplanin knock-out (PDPN KO; blue) B16F10 murine melanoma cell lines (right). mRNA expression is calculated as fold change and normalized to *Pdgfra* (LN stromal cells) or *Gapdh* (B16F10) expression. Data shown as mean with dots representing n=6 (LN stromal cells) or n=3 (B16F10) biological replicates. **B**. Analysis of podoplanin surface expression in PDPN^+^ (red) and PDPN KO (blue) B16F10 cell lines by flow cytometry. Cells not stained with antibody (no Ab; grey) are used as negative control. gMFI = geometric mean fluorescence intensity. **C**. Left: Immunofluorescence of F-actin (white) in PDPN^+^ (top) and PDPN KO (bottom) B16F10 cell lines, labelled with mOrange (red) or CFP (blue) respectively. Maximum Z stack projections of representative images from n=3 biological replicates are shown. The scale bars represent 100 microns. Right: Cell area (in *µ*m^2^; top) and perimeter (in *µ*m; bottom) of PDPN^+^ (red) and PDPN KO (blue) B16F10 cells. Dots represent single cells. n=77-90 cells collated from 3 biological replicates. Error bars represent median with interquartile range. \*\*\*\**p*<0.0001. **D**. Left: Immunofluorescence imaging of F-actin (white) in mixed PDPN^+^ (red) and PDPN KO (blue) B16F10 tumour 9 days post-injection. The scale bars represent 100 (top) or 50 (bottom; zoom) microns. Right: Cell area of PDPN^+^ (red) and PDPN KO (blue) B16F10 cells in the tumour. Dots represent single cells (n=97-197 cells). Error bars represent median with interquartile range. \*\*\*\**p*<0.0001.

We then asked whether podoplanin-driven contractility can control melanoma cell morphology in the tumour microenvironment *in vivo*. All tumour areas contain both PDPN^+^ and PDPN KO cells in varying proportions suggesting some degree of migratory behaviour *in vivo*, or tissue fluidity, resulting in cell mixing as opposed to clonal segregation. Podoplanin expression is maintained in PDPN^+^ cells *in vivo* (**Fig. S2**). PDPN^+^ cells have a smaller cross-sectional area than PDPN KO cells in the same tumour regions (**Fig. 2D**), and moreover, cortical F-actin structures are also more prominent in PDPN^+^ cells (**Fig. 2D**). These results are consistent with the hypothesis that podoplanin intrinsically drives contractility of the actin cytoskeleton in melanoma cells to result in a rounded amoeboid-like cell morphology.

### Increased cell motility and invasion of podoplanin^+^ melanoma cells *in vivo*

Upon examination of the cellular ratios of PDPN^+^ (mOrange) vs PDPN KO (CFP) at the tumour boundary of mixed tumours, we observed that PDPN^+^ B16F10 cells are consistently overrepresented in the majority of tumour margins (5/7 tumour sections), when controlling for the varying ratio of PDPN^+^:PDPN KO throughout the whole tumour section (**Fig. 3A** and **Fig. S3**). Beyond the tumour boundary, the tumour capsule is frequently populated by host PDPN^hi^ cells likely to be fibroblasts. Within the fibroblastic-rich tumour capsule, the locally invasive tumour cells are almost exclusively PDPN^+^ cells (**Fig. 3B**). These invasive PDPN^+^ tumour cells have a rounded morphology and are exclusively observed invading as single cells (**Fig. 3B**). Local invasion of individual cancer cells through the matrix requires a balance of proteolysis, and high cytoskeletal contractility permitting amoeboid invasion. The proteolytic activity of PDPN^+^ and PDPN KO melanoma cells is comparable (**Fig. S4**) indicating that podoplanin is not promoting local invasion via increased proteolysis in this model. We next sought to ask whether overexpression of podoplanin (PDPN-CFP) in PDPN^lo^ 5555 melanoma cells (**Fig. S1A**) is sufficient to enhance amoeboid motility *in vivo*. We expressed PDPN-CFP in a subset of mCherry-labelled 5555 cells. The 5555-mCherry cells which overexpress PDPN-CFP move as single cells with largely rounded morphology and rapid changing cell shape (**Supplementary Movie 1** and **Fig. 3C**). 63% of 5555-mCherry cells overexpressing PDPN-CFP are motile or migrating compared to 51% of total 5555-mCherry cells *in vivo* (**Fig. 3D**). This indicates that overexpression of podoplanin increases amoeboid motility *in vivo*.

**Fig. 3.**
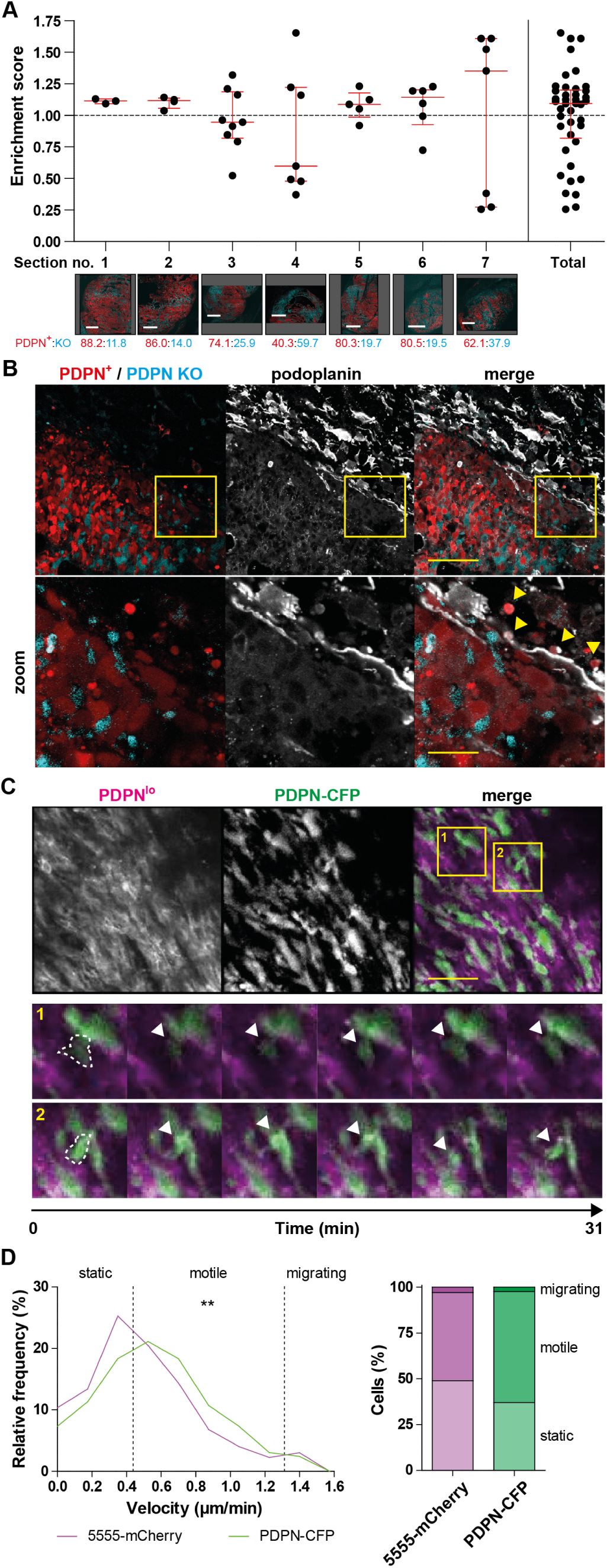
Podoplanin dependent transition to amoeboid dissemination in melanoma cells. **A**. Enrichment of PDPN^+^ B16F10 cells in the invasive front (IF) compared to the total number of cells in the whole mixed PDPN^+^/PDPN KO tumour. Enrichment score above 1 (dashed line) indicates relative enrichment of PDPN^+^ cells in the IF. 7 individual sections of 2 different tumours were analysed using QuPath software (**Fig. S3**). Data shown as median with interquartile range. Tilescans of each tumour section are depicted below the graph with the respective ratio of PDPN^+^ vs. PDPN KO areas. **B**. Immunofluorescence imaging of podoplanin (white) in mixed PDPN^+^ (red) and PDPN KO (blue) B16F10 tumour 9 days post-injection. Arrow heads indicate disseminated PDPN^+^ B16F10 cells. The scale bars represent 100 (top) or 50 (bottom; zoom) microns. **C**. Time-lapse imaging of mixed PDPN^lo^ (mCherry-labelled; magenta) and PDPN-CFP (green) transfected 5555 murine melanoma tumour (see also Supplementary Movie 1). Dashed line indicates PDPN-CFP^+^ cell tracked over time (arrow heads). The scale bar represents 30 microns. **D**. Left: Velocity of PDPN^lo^ (mCherry-labelled; magenta) and PDPN-CFP (green) transfected 5555 cells *in vivo*. \*\**p*=0.0013. Right: Percentage of static, motile or migratory PDPN^lo^ (mCherry-labelled; magenta) and PDPN-CFP (green) transfected 5555 cells. **Supplementary Movie 1**. Time-lapse imaging of mixed PDPN^lo^ (mCherry-labelled; magenta) and PDPN-CFP (green) transfected 5555 murine melanoma tumour. Stars in first frame indicated the example PDPN-CFP+ cells shown in **Fig. 3C**.

### Podoplanin expression drives dedifferentiation of melanoma

We observed that mixed tumours of PDPN^+^/PDPN KO are often predominantly PDPN^+^ despite transferring equal cell numbers (**Fig. S5A**). Neither podoplanin expression nor the fluorescence protein expressed had any effect on cell proliferation *in vitro* (**Fig. S5B**). However, we do find that PDPN^+^ tumours grow much more rapidly *in vivo* than PDPN KO tumours (**Fig. S5C**), suggesting that PDPN expression confers some survival advantage.

Acquisition of an invasive motile phenotype is linked with dedifferentiation of melanoma, marked by loss of melanocyte functions such as pigmentation (9). We observe that PDPN KO B16F10 tumours are more pigmented compared to PDPN^+^ tumours (**Fig. 4A**). This altered pigmentation is cell intrinsic and consistently maintained in mixed *in vitro* cultures (**Fig. 4B**), suggesting that knock-out of podoplanin expression restores pigmentation, a characteristic of non-invasive and non-cancerous melanocytes (9).

**Fig. 4.**
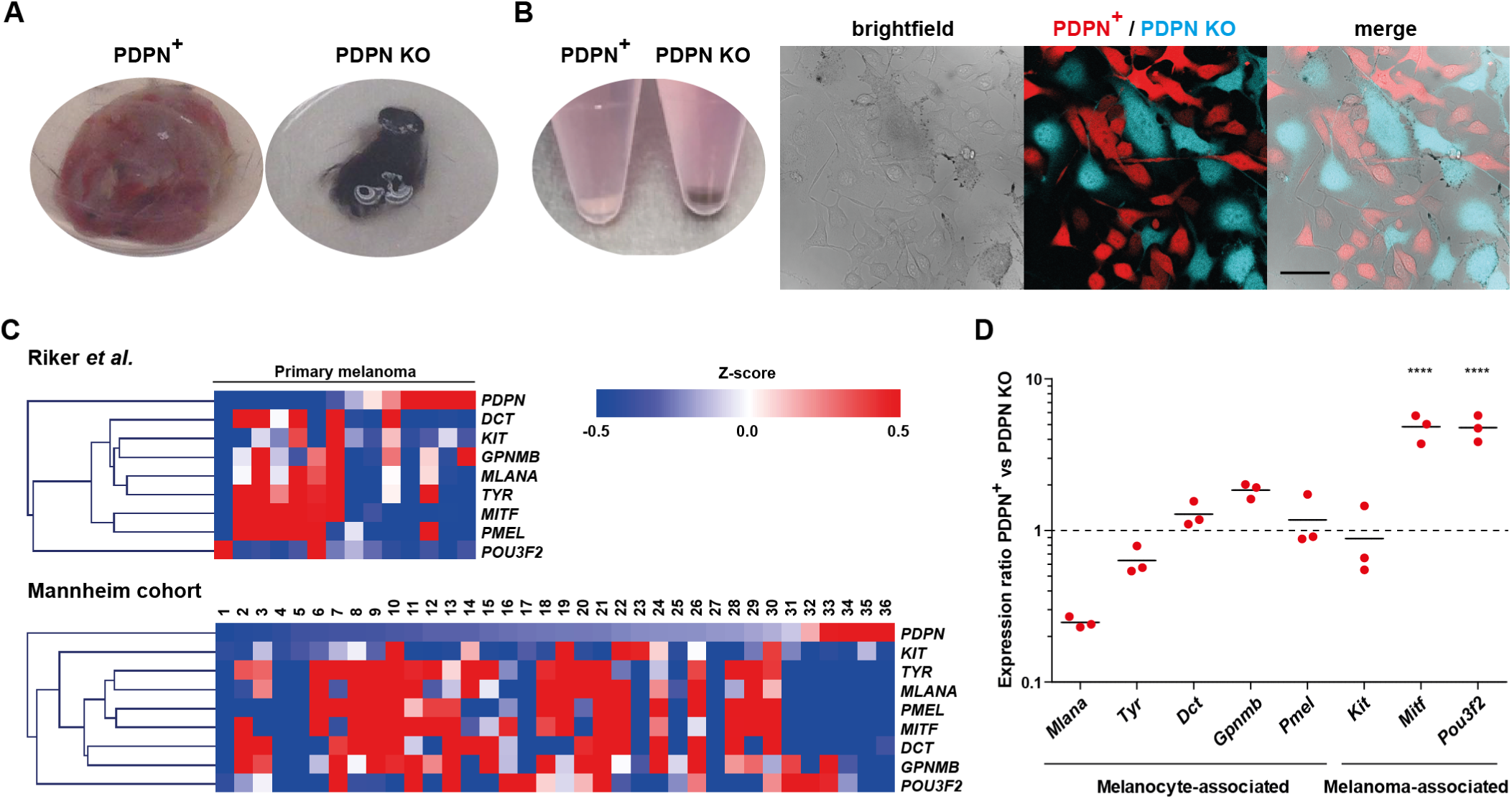
Loss of podoplanin restores pigmentation and melanocyte differentiation. **A**. Tumours (left) and cells pellets (right) of PDPN^+^ and PDPN KO B16F10 cell lines. **B**. Imaging of pigmentation (brightfield; left) of PDPN^+^ and PDPN KO B16F10 cell lines, labelled with mOrange (red) or CFP (blue) respectively (middle). The scale bar represent 50 microns. **C**. Heatmaps showing expression (Z-score) of podoplanin (*PDPN*) and eight dedifferentiation-associated genes in datasets of primary tumour samples of melanoma patients (top; Riker *et al*. GSE7553) and metastatic melanoma cultures (bottom; Mannheim cohort, GSE4843; from Hoek *et al*. 2006). For the Mannheim cohort, each number indicates a separate metastatic melanoma culture. **D**. mRNA expression of five melanocyte-associated (*Mlana, Tyr, Dct, Gpnmb, Pmel*; left) and three invasion-associated (*Kit, Mitf, Pou3f2*; right) genes in PDPN^+^ B16F10 cells. mRNA expression is calculated as fold change of *Gapdh* expression and normalized to expression in PDPN KO B16F10 cells (set at 1 as indicated by dashed line). Data shown as mean with dots representing n=3 biological replicates. \*\*\*\**p*<0.0001.

To explore whether podoplanin expression altered gene expression critical to pigmentation, we examined mRNA expression in human primary melanoma patient samples and cell lines. When clustered based on *PDPN* expression, we found that the *PDPN*^hi^ samples expressed lower levels of key melanocyte genes including tyrosinse (*TYR*) and pre-melanosome protein (*PMEL*) (**Fig. 4C**), providing a mechanistic explanation for the changes in pigmentation we observe in PDPN^+^ vs PDPN KO melanoma xenograft tumours and cell lines (**Fig. 4A-B**). We therefore directly tested the impact of podoplanin on melanocyte gene expression. Melanocyte-associated markers required for pigmentation, *Mlana, Pmel* and *Tyr* (9), are also downregulated in PDPN^+^ B16F10 mouse melanoma cell line compared to PDPN KO controls, whereas the melanoma-associated genes *Mitf* and *Pou3f2* are upregulated (**Fig. 4D**). *Pou3f2* encodes the transcription factor Brn2, a major regulator of melanoma phenotype switching, and directly linked with dedifferentiation and acquisition of motility (8, 10). These data provide evidence that high podoplanin expression identifies a more dedifferentiated subset of melanoma, linked to a switch towards a more malignant and invasive phenotype. Since high podoplanin-expressing melanoma downregulates melanocyte markers, it may be harder to identify these aggressive tumours using standard diagnostic biomarkers.

## Discussion

It is known that cancer cells can utilise amoeboid migration for local invasion and initiation of metastasis, and the cytoskeletal rearrangements controlling this mode of migration are well studied. However, how these signalling pathways become hyperactivated are not fully understood, and we lack druggable molecular targets to inhibit amoeboid invasion. Here, we show that expression of podoplanin, a known driver of actomyosin contractility in fibroblasts, is upregulated in a subset of melanoma patients. We show that podoplanin is able to drive high cytoskeletal contractility in melanoma cells promoting an invasive amoeboid phenotype. In agreement with our presented data, high podoplanin expression has been observed at the invasive front of different cancer types (13), where individual invading amoeboid cells are also observed (11, 14, 15).

Pro-inflammatory cytokines have been linked to increased actomyosin contractility and amoeboid invasion in stroma and tumour cells (11, 16). Additionally, cytokine signalling in the tumour microenvironment is one pathway known to induce aberrant expression of podoplanin (13). Podoplanin expression can also be induced by the activity of oncogenes such as Src kinase (17), and furthermore the transcription factor AP-1, which directly binds the podoplanin promotor in malignant keratinocytes resulting in increased podoplanin expression in skin tumours (18). We have recently shown that binding of the podoplanin binding partner, C-type lectin-like receptor 2 (CLEC-2) in addition to inhibiting podoplanin-driven contractility (12), also upregulates podoplanin expression (19). CLEC-2 is highly expressed on platelets, which may leak into the tumour via poorly functioning blood vessels. This interaction may further upregulate podoplanin expression on tumour cells, leading to more amoeboid and invasive phenotype.

It was already known that podoplanin expression was correlated with poor prognosis and higher incidence of metastatic disease (13), but the mechanisms were not fully understood. We now show that podoplanin is a driver of amoeboid invasion, and furthermore we link podoplanin directly to melanoma dedifferentiation. The amoeboid cell state supports both invasion and tumour cell ‘stemness’ (20). Our data supports that podoplanin expression drives a wide-ranging re-programming of melanoma towards a de-differentiated, invasive phenotype with higher capacity for tumour initiation or proliferation. We predict that targeting podoplanin to block downstream signalling in melanoma would reverse dedifferentiation, and as such inhibit invasion and metastasis. A potential strategy could be to design small-molecules to mimick CLEC-2 binding to podoplanin. Podoplanin has potential as a diagnostic biomarker for dedifferentiated melanoma, as well as migrastatic drug target, which could be used in combination with current treatments for melanoma.

## Materials and Methods

### Mice

Wild-type C57BL/6J mice were purchased from Charles River Laboratories. Both males and females were used for *in vivo* experiments and were aged 6-10 weeks. All mice were age-matched and housed in specific pathogen-free conditions. All animal experiments were reviewed and approved by the Animal and Ethical Review Board (AWERB) within University College London and approved by the UK Home Office in accordance with the Animals (Scientific Procedures) Act 1986 and the ARRIVE guidelines.

### Cell culture

WM983A (primary) and WM983B (invasive/metastatic) human melanoma cell lines are previously described (14). Murine melanoma cell lines 4434 and 5555 (kind gift from Prof. Dr. R. Marais, The University of Manchester, Manchester, UK), and B16F10 (kind gift from Prof. Dr. C. Reis e Sousa, The Francis Crick Institute, London, UK) are all originally generated in C57BL/6 mice. B16F10 cell lines stably expressing mOrange or CFP were generated using the piggyBac transposon-based expression system. B16F10 cells were transfected with pBSR2-mOrange or pBSR2-CFP, and pBase plasmids (kind gift from Dr. E. Sahai, The Francis Crick Institute, London, UK). 5555 cell lines were labeled with mCherry membrane marker (GAP43-mCherry (kind gift from Dr. E. Sahai, The Francis Crick Institute, London, UK)) for intravital imaging. Additionally, PDPN-CFP (12) was overexpressed for gain-of-function experiments.

Stable podoplanin knock-out (PDPN KO) B16F10 cell lines were generated using CRISPR/Cas9 editing. B16F10 cell lines were transfected with pRP[CRISPR]-hCas9-U6>PDPN gRNA 1 plasmid (constructed and packaged by Vector-builder; vector ID: VB160517-1061kpr) before negative selection using MACS LD columns (Miltenyi Biotec), anti-mouse podoplanin-biotin antibody (50 *µ*g/mL, clone 8.1.1, eBioscience, 13-5381-82), and anti-biotin microbeads (Miltenyi Biotec, 130-090-485).

Primary murine lymph node stromal cells were isolated and cultured for 3 days *ex vivo* as previously described (12). Murine and human melanoma cell lines were cultured in DMEM with GlutaMAX (Gibco, via Thermo Fisher Scientific) supplemented with 10% fetal bovine serum (Sigma-Aldrich) and 100U/mL penicillin-streptomycin (Gibco, via Thermo Fisher Scientific) at 37°C, 5% CO_2_ (B16F10) or 10% CO_2_, and passaged using Trypsin/dPBS (Gibco, via Thermo Fisher Scientific).

### *In vitro* cell proliferation assay

mOrange or CFP-labelled PDPN^+^ or PDPN KO B16F10 cells (5×10^4^ cells per well) were seeded in individual wells in 6-well plates (1 plate per time point). Every 24 hours, cells were harvested and diluted 1:2 in 0.4% Trypan Blue Solution (Thermo Fisher Scientific), and counted using a Neubauer Haemocytometer Counting Chamber (0.1mm; Camlab).

### *In vitro* proteolysis assay

12-well glass-bottomed culture plate (MatTek) was coated with Oregon Green 488-conjugated gelatin (Invitrogen; diluted 1:8 in solution of 2.5% (w/w) gelatin/2.5% (w/w) sucrose (Sigma-Aldrich)) for 10min at room temperature (RT). B16F10 cells were seeded (1.2×10^4^ cells per well) and incubated at 37^*°*^C, 5% CO_2_. After 20 hours, cell cultures were fixed with 3.6% formaldehyde (Sigma-Aldrich; diluted in PBS), and permeabilized with 0.2% Triton-X100 (Sigma-Aldrich) in PBS for 15min at RT. Cell nuclei were visualized using Hoechst (1:12000 dilution, Fisher Scientific) by 1h incubation at RT. Stained samples were stored in PBS at 4^*°*^C until imaging.

### *In vivo* tumour growth

mOrange-labelled PDPN^+^ and/or CFP-labelled PDPN KO B16F10 cells were injected subcutaneously into the flank of wild-type C57BL/6J mice (total 1×10^6^ cells in 100 *µ*l PBS per mouse). Tumours were excised 9-10 days after injection, and individual tumours were weighed. Tumours were fixed in AntigenFix (DiaPath, via Solmedia) overnight (o/n) at 4^*°*^C, and subsequently transferred to 30% sucrose for o/n incubation at 4^*°*^C. Tumours were frozen in O.C.T Compound (Tissue-Tek) using dry ice and iso-pentene, and stored at -80^*°*^C.

### Intravital imaging

C57BL/6J mice were injected subcutaneously with 1:1 mix of PDPN^lo^ 5555 (mCherry) and PDPN-CFP 5555. When tumours were 3-7mm diameter, mice were anaesthetised using isofluorane and a skin flap was cut to expose the tumour before the mouse was positioned on an Zeiss880 laser scanning microscope connected to a Chameleon Coherent Ti-Sapphire laser tuned to 850 nm. Anesthesia was maintained while time-lapse movies were made of the tumours. Each region was imaged for 20-30 minutes for each tumour. Following acquisition, timelapse stacks were drift corrected and using Imaris, movies are presented as z-projections. Analysis of cell movement was performed using the spots function in Imaris software.

### Immunofluorescence imaging and analysis

For analysis of cell morphology of WM983 cell lines, 2×10^4^ cells were seeded on top of 1.7mg/mL collagen (type I, rat tail) with or without Matrigel matrix (both from Corning, via Thermo Fisher Scientific) supplemented with 10% minimum essential medium alpha medium (MEMalpha, Invitrogen, via Thermo Fisher Scientific) and 10% FCS (Greiner Bio-One). For analysis of F-actin organisation in B16F10 cells *in vitro*, 3×10^4^ cells were seeded in 35mm glass-bottomed culture dishes (MatTek) at 37^*°*^C, 10% CO_2_. At 24h, cell cultures were fixed with AntigenFix (DiaPath, via Solmedia) for 15min at RT or 3.6% formaldehyde (Sigma-Aldrich; diluted in PBS), and permeabilized with 0.2% Triton-X100 (Sigma-Aldrich) in PBS for 15min at RT. F-actin and cell nuclei were visualized using respectively phalloidin-TRITC (P1951-1MG) and DAPI (D9542-1MG; both 1:500 dilution, both from Sigma-Aldrich).

B16F10 tumours were cyrosectioned at 20 *µ*m. Frozen tumour sections were air dried for 10min at RT, encircled with a PAP pen for immunostaining (Sigma-Aldrich) and fixed with AntigenFix (DiaPath, via Solmedia) for 20min at RT. Tissue sections were permeabilised and blocked with 0.3% Triton, 1% mouse serum and 2% BSA in 0.1M Tris-HCL buffer (pH 7.4) for 1h, and stained with hamster anti-mouse podoplanin (1:500, clone 8.1.1, DM3501, Acris Antibodies) o/n at 4^*°*^C, then incubated with goat-anti-hamster IgG AlexaFluor 647 (1:500; Invitrogen, via Thermo Fisher Scientific) and phalloidin-TRITC (1:500; P1951-1MG) for 2h at RT, and mounted in Mowiol (Sigma-Aldrich).

All immunofluorescence images were acquired using a Leica DMI6000 SP5 confocal microscope using HC PL FLUOTAR /0.3 10x air (WM983 cell lines), or HCX PL APO /1.25 40x or /1.4 63x oil (B16F10 cell lines and tumour sections) objective lenses.

Immunofluorescence images were analysed using Fiji/ImageJ software. Z stacks were projected with ImageJ Z Project (maximum projection). Roundness index, cell area and perimeter were analysed by manually drawing around the cell shape using F-actin staining, or mOrange or CFP-labelling.

Tilescans of tumour sections taken at various depths of mixed PDPN^+^/KO tumours from 2 mice were analysed using QuPath 0.2.0 (**Fig. S3**). Using the pixel classifier tool, the software was trained based on defined PDPN^+^ (mOrange), PDPN KO (CFP) and tumour-free regions. A pixel classification was then created and applied to all images, dividing sections into PDPN^+^, PDPN KO and tumour-free areas. Annotations were created around the whole tumour section, and overall proportions of PDPN^+^:PDPN KO were automatically generated. Invasive front sections were defined as 30 *µ*m from the tumour edge. The margin was classified where the bulk of tumour fluorescence ended. Enrichment scores were calculated by normalizing the proportion of PDPN^+^ areas at invasive edges to overall PDPN^+^ areas in the whole tumour section.

### Flow cytometry

Single-cell suspensions of murine melanoma cell lines were incubated with FcR blocking reagent (Miltenyi Biotec) as per supplier’s instructions, followed by staining with hamster anti-mouse podoplanin-eFluor660 antibody (1:200, clone 8.1.1, eBioscience, 50-5381-82) diluted in PBS supplemented with 0.5% BSA and 5mM EDTA for 30min on ice. Stained cells were analysed using FACSDiva software and LSRII flow cytometer (BD Biosciences). Flow cytometry data was analysed using FlowJo Software version 10 (BD Biosciences).

### RNA isolation and quantitative RT-PCR analysis

mRNA of primary lymph node stromal cells, WM983, and B16F10 cell lines was isolated using RNeasy Mini kit (Qiagen) as per supplier’s instructions, including a DNA digestion step. Then reverse transcribed to cDNA using SuperScript IV First-Strand Synthesis System kit (Invitrogen, via Thermo Fisher Scientific) as per supplier’s instructions. Transcript abundance of genes of interest were determined with a CFX96 Sequence Detection System (Bio-Rad) with MESA Blue (Eurogentec, via Promega). Gene-specific oligonucleotide primers are listed in Table 1 (see Supplementary Information). Ct values of the genes of interest were normalized to the Ct value of the housekeeping gene *GAPDH* (WM983 cell lines), *Pdgfra* (lymph node stromal cells) or *Gapdh* (B16F10 cell lines).

**Table 1.**
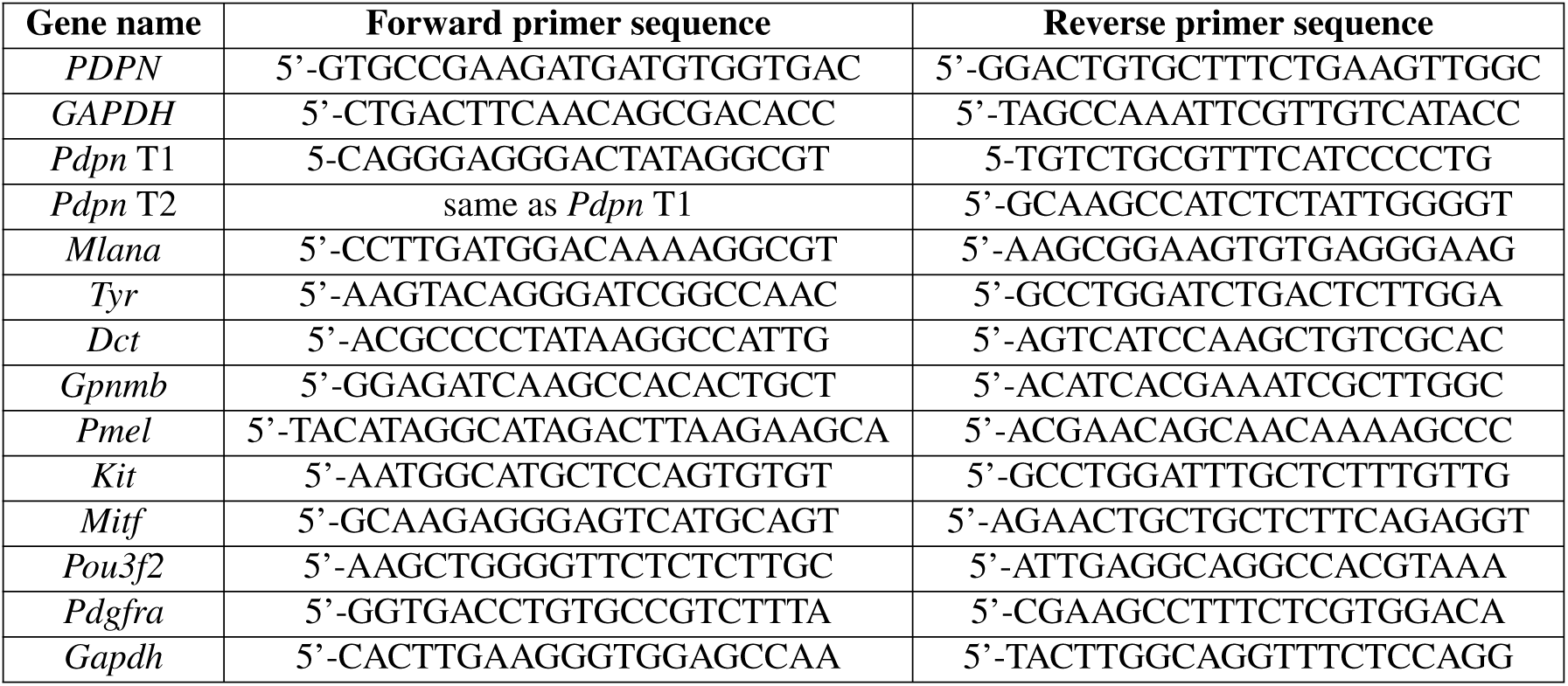
Gene-specific oligonucleotide primers used for quantitative RT-PCR analysis

### Melanoma patient and cell lines mRNA database analysis

Expression data from melanoma patient public studies Kabbarah *et al* (GSE46517) and Riker *et al* (GSE7553) were extracted from GEO (Gene Expression Omnibus) database and normalized using Gene Pattern software (http://www.broadinstitute.org/cancer/software/genepattern/). Expression data from melanoma cell lines (Mannheim cohort; GSE4843) was obtained from the study Hoek *et al*, 2006. One outlier was excluded from the Mannheim cohort.

### Statistics

Statistical differences between two groups were determined using unpaired Student’s t-test (two-tailed), or, in the case of non-Gaussian distribution, Mann-Whitney test. Statistical differences between expression of melanocyte and invasion-associated genes in PDPN^+^ and PDPN KO B16F10 cells were determined using two-way ANOVA with Sidak’s multiple comparisons test. Statistical tests were performed using GraphPad Prism software (version 7), and differences were considered to be statistically significant at *p*≤ 0.05.

## ACKNOWLEDGEMENTS

This work is funded by Cancer Research UK Career development fellowship CRUK-A19763 (to S.E.A.), C33043/A12065 and C33043/A24478 (to V.S.-M.), Rubicon Postdoctoral fellowship from the Dutch Research Council (NWO) 019.162LW.004 (to C.M.d.W.), Barts Charity and Medical Research Council (MC-U12266B).

---

**Fig. S1.**
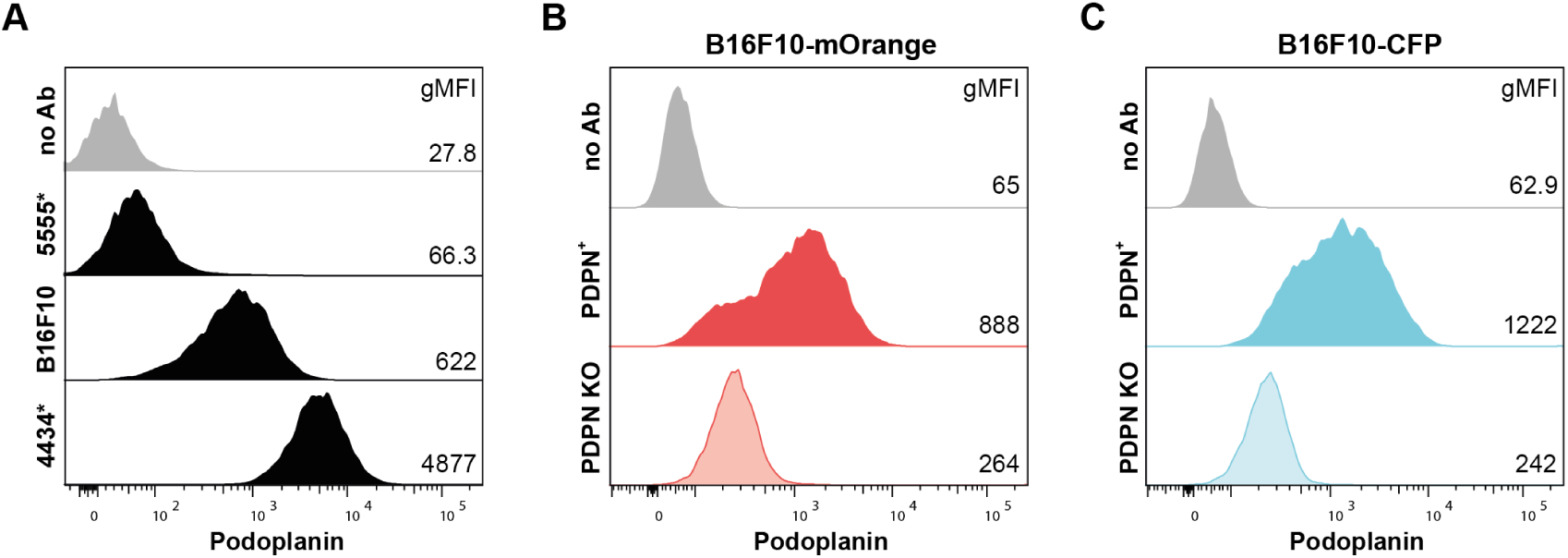
**A**. Analysis of podoplanin surface expression in three murine melanoma cell lines (from top to bottom 5555, B16F10 and 4434), **B**. mOrange-labelled and **C**. CFP-labelled PDPN+ (solid histograms) and PDPN KO (tinted histograms) B16F10 cell lines by flow cytometry. Cells not stained with antibody (no Ab; grey) are used as negative control. gMFI = geometric mean fluorescence intensity. * = 5555 and 4434 cell lines are *Braf* -V600E mutation positive.

**Fig. S2.**
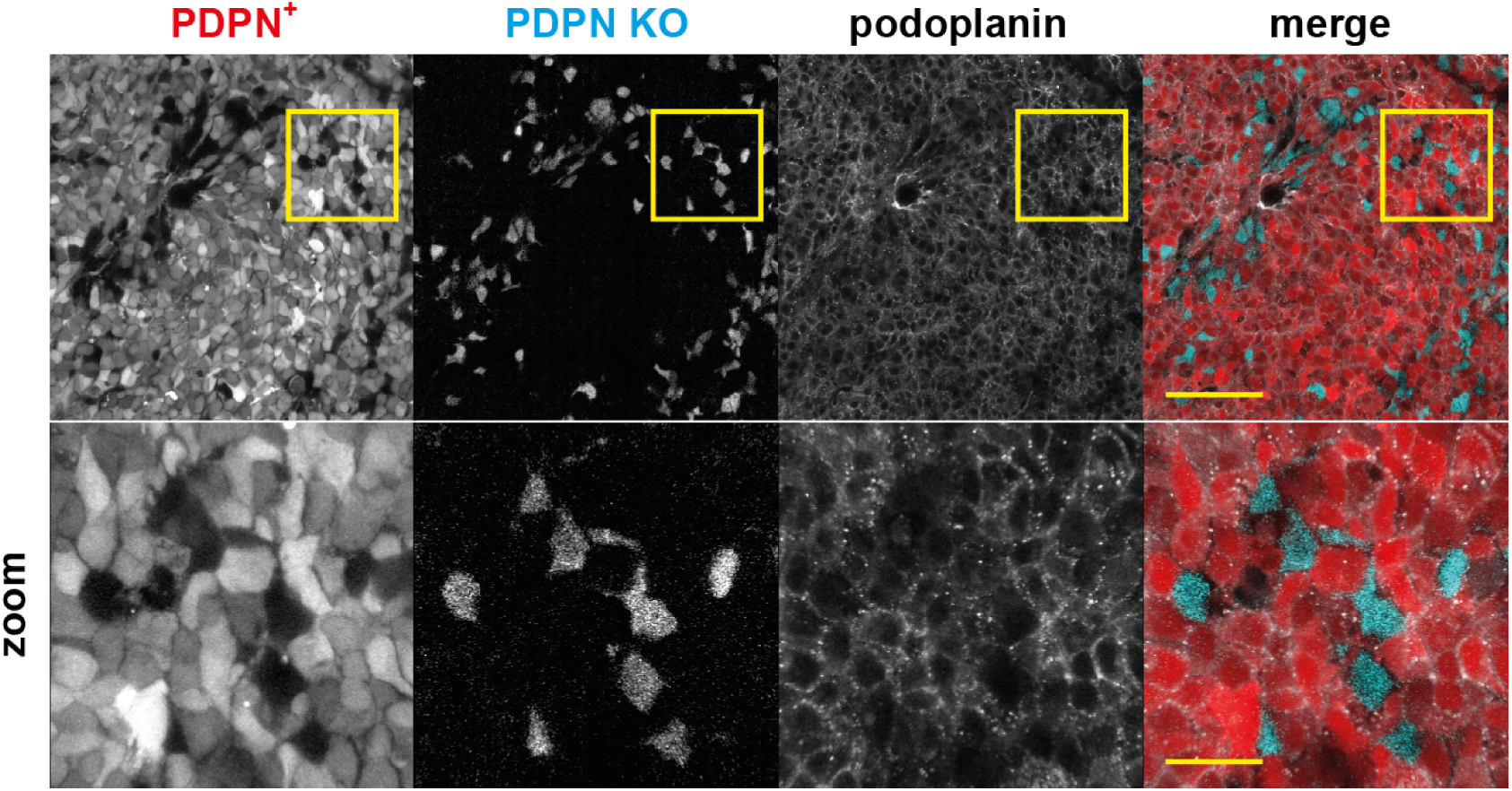
Immunofluorescence imaging of podoplanin (white) in mixed PDPN+ (red) and PDPN KO (blue) B16F10 tumour 9 days post-injection. The scale bars represent 100 (top) or 50 (bottom; zoom) microns.

**Fig. S3.**
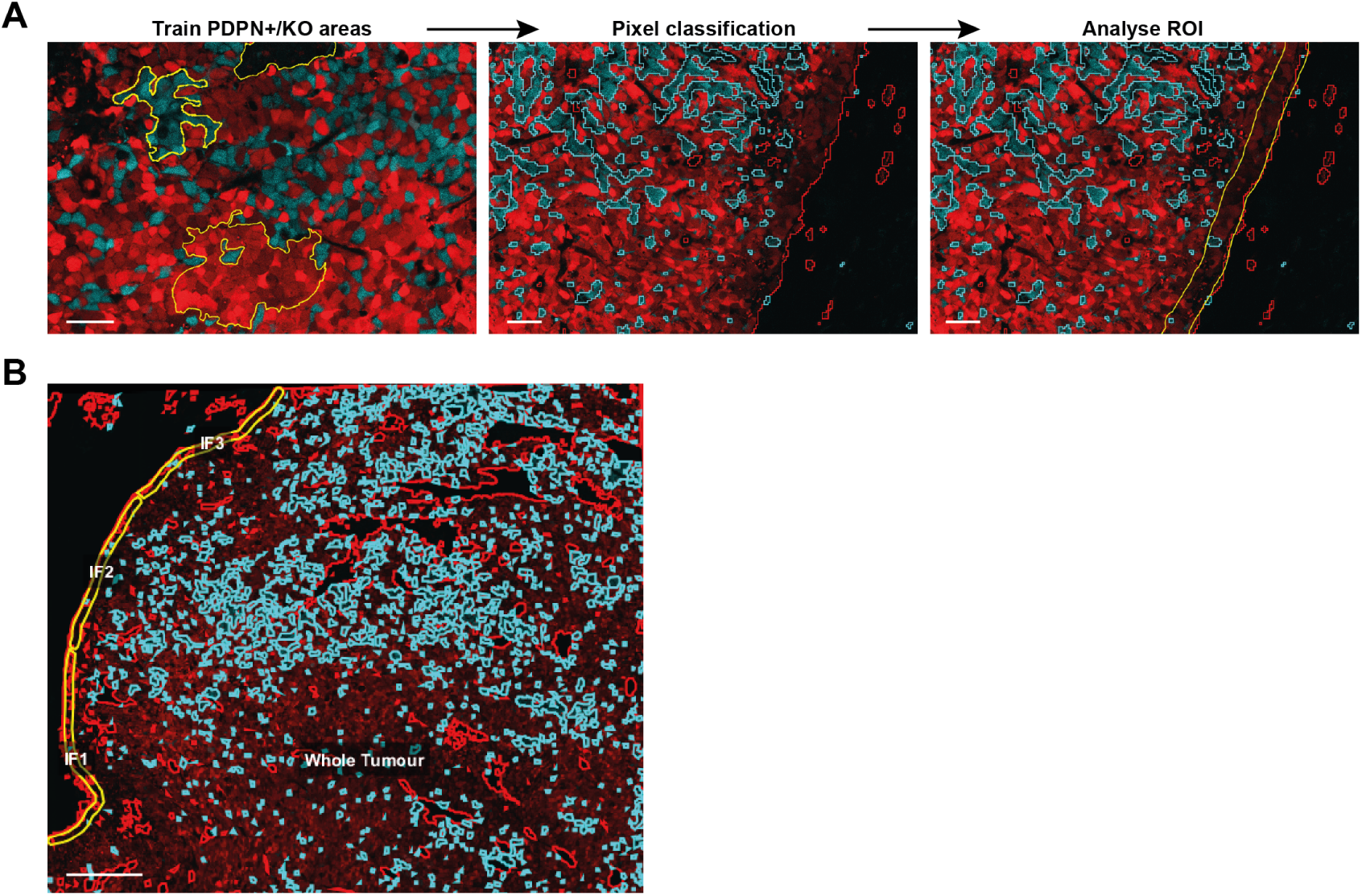
Analysis of mixed B16F10 PDPN+ (red) and PDPN KO (cyan) tumour sections using QuPath software. **A**. Workflow. PDPN+, PDPN KO and tumour-free areas were annotated using the wand tool and allocated into specific classes to train the software (left). Pixel classification was created and applied to all images, separating sections into PDPN+, PDPN KO or tumour-free areas (middle). Regions of interest (ROI) of the whole tissue section and invasive front at 30 microns (yellow) are drawn, and proportion of PDPN+ and PDPN KO cells are calculated automatically (right). The scale bars represent 50 microns. **B**. Tissue section of mixed PDPN+/KO B16F10 tumour analysed by QuPath workflow as shown in A. For each tumour section, multiple invasive front (IF; yellow) areas were identified around the tumour edge, and the proportion of PDPN+ vs PDPN KO cells were calculated for each IF. The scale bar represents 250 microns.

**Fig. S4.**
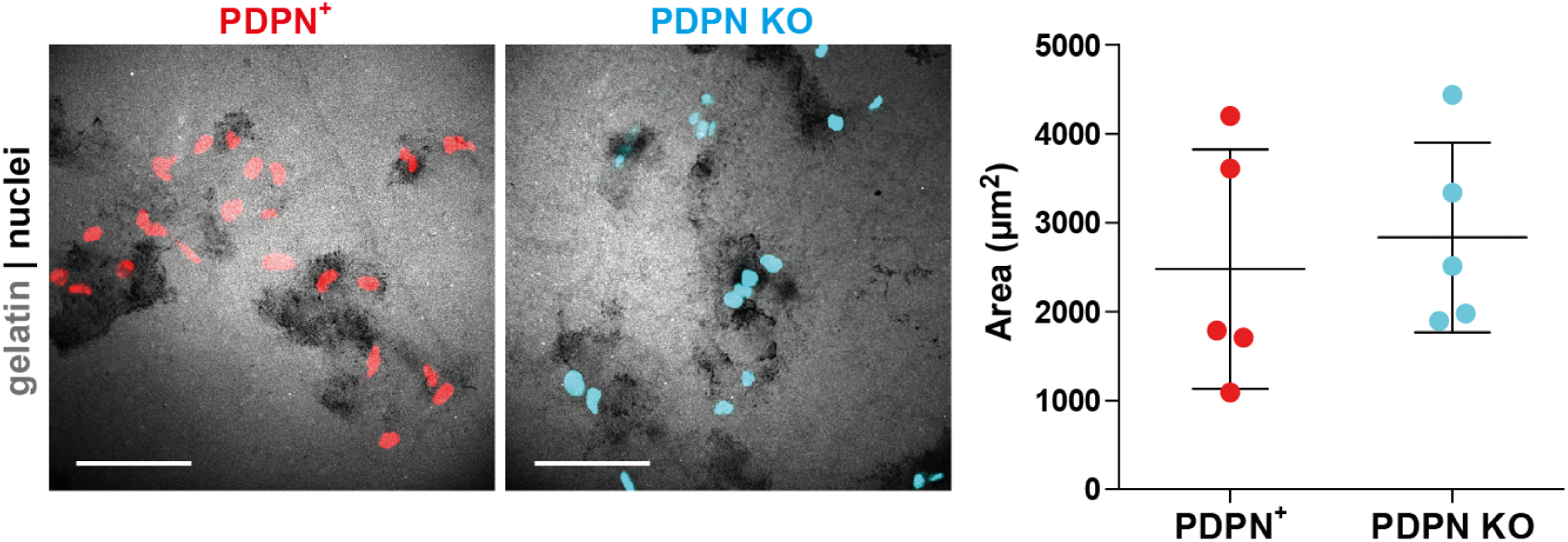
*In vitro* proteolysis of gelatin by PDPN+ and PDPN KO B16F10 cell lines. Cell nuclei are visualized with Hoechst (red for PDPN+ nuclei, cyan for PDPN KO nuclei). The scale bars represent 100 microns. Right: Area in *µ*m^2^ of gelatin degradation normalized for number of PDPN+ (red) and PDPN KO (cyan) B16F10 cells per field of view. Data shown as mean +/-SD with dots representing individual fields of view.

**Fig. S5.**
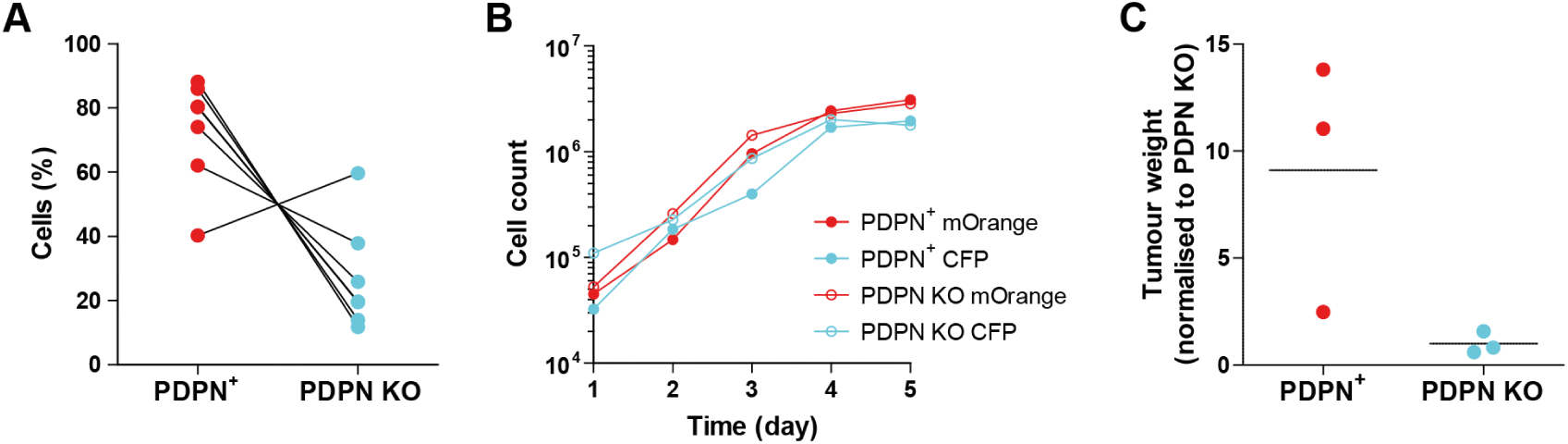
**A**. Percentage of PDPN+ and PDPN KO B16F10 cells in individual sections of mixed tumours as shown in Fig. 3A. **B**. *In vitro* proliferation of mOrange-labelled (red) and CFP-labelled (blue) PDPN+ (closed circles) and PDPN KO (open circles) B16F10 cell lines during 5 days. **C**. Weight of dissected PDPN+ (red) and PDPN KO (blue) B16F10 tumours 10 days post-injection. Data is normalized to average weight of PDPN KO B16F10 tumours (set at 1), and shown as mean with dots representing individual tumours.

